# Evaluating an Upper Respiratory Disease Panel on the Portable MinION Sequencer

**DOI:** 10.1101/436600

**Authors:** Wanda J. Lyon, Zachary K. Smith, Brian Grier, James Baldwin, Clarise R. Starr

**Affiliations:** 711^th^ Human Performance Wing, RHXB, Wright Patterson Air Force Base OH, USA; 711^th^ Human Performance Wing, USAFSAM, Wright Patterson Air Force Base, OH, USA

## Abstract

The MinION was used to evaluate upper respiratory disease infections using both whole genome amplification (WGA), targeted sequencing, and was found to have tremendous potential for field use. The MinION nanopore sequencer was been released to community testers for evaluation using a variety of sequencing applications. The MinION was used to evaluate upper respiratory disease infections using both whole genome amplification and targeted sequencing, and was found to have tremendous potential for field use. In this study, we tested the ability of the MinION nanopore sequencer to accurately identify and differentiate clinical bacterial and viral samples via targeted sequencing and whole genome sequencing. The current nanopore technology has limitations with respect to error rate but has steadily improved with development of new flow cells and kits. Upper respiratory disease organisms were successfully identified and differentiated down to the strain level with 87-98% alignment to our reference genome database. The ability to differentiate strains by amplicon and whole genome sequencing on the MinION was accomplished despite the observed average per 100-base error rate averaged 1.2E-01. This study offers evidence of the utility of sequencing to identify and differentiate both viral and bacterial species present within clinical samples.

## Introduction

Acute respiratory infections (ARI) have been a significant source of disease and nonbattle injury among military forces for centuries. Although mortality from ARIs is low in military populations, the impact of these infections on military readiness in terms of lost work-hours and convalescence is high. Furthermore, novel ARIs have been a periodic source of great morbidity and mortality, such as during the 1918 A/H1N1 (“Spanish”) influenza epidemic, the 2002-2003 severe acute respiratory syndrome (SARS) epidemic in Asia and North America, and the 2009-2010 swine-origin A/H1N1 global pandemic. Periodic outbreaks of adenovirus and similar infections have plagued military training centers in recent years, control of which has been hampered until very recently by the unavailability of an active vaccine [12, 17]. Based on historical information dating back to the 1918 influenza pandemic, ARI has also had a major impact on military training and readiness. Supporting the notion that viral infections play a major role in training and readiness, other studies have demonstrated rates of ARI in 80% of trainees, with rates of hospitalization of 20% prior to influenza and adenovirus vaccination programs [15]. Despite aggressive vaccination programs, ARI has been noted to be the cause of 70% of U.S. Air Force pilot groundings and led to sick call evaluations in 1.5% of all military members seeking care during Operation Desert Storm [24]. Nonvaccine preventable causes of ARI in military members include respiratory syncytial virus, enteroviruses, and rhinoviruses [14, 32]. Although influenza is usually the pathogen that captures the attention of the world, other respiratory pathogens have recently garnered the attention of the military. Adenoviruses cause a wide array of symptoms, from conjunctivitis, gastroenteritis, and mild respiratory infections to life-threatening pneumonias in young adults and blood stream and brain infections in immunocompromised hosts. Enterovirus 68 has recently shown severe respiratory illness in other countries such as the Philippines, the Netherlands, and Japan [13, 19], and the Middle East respiratory syndrome-coronavirus (MERS-CoV) outbreak in the Middle East [33, 34] along with the H7N9 influenza outbreak in China [39] has put focus on new and emerging pathogens that may be introduced into immunologically naïve populations or recombined into novel forms through human-human or human-animal transmission.

Next-generation sequencing (NGS) technologies are now capable of providing a whole genome sequence or metagenomics for a wide range of organisms [16, 36]. NGS is being applied with increasing frequency in clinical microbiology laboratories to detect and characterize pathogens, thereby replacing the currently used Sanger sequencing methods [3, 39]. Sanger sequencing has been a reliable and robust method that has served clinical laboratories well for over several decades; however, it is labor intensive, slow, and not easily adapted for processing large genomes or large samples sets.

Oxford Nanopore’s MinION may therefore provide us with new opportunities to track infectious disease, for example, rapid sequencing of viral genomes in response to the early phase of influenza pandemics, or for determination of coronavirus virus genotypes during outbreaks. In addition, sequencing will allow for rapid sequencing of viral genomes and provides an opportunity to gain insight into viral genetic drift (single nucleotide polymorphism), emergence of new strains such as SARS-like coronavirus or MERS-CoV, and transmission. To date, a limited number of pathogenic viral studies have been done with the MinION, and all of the published results have focused on sequencing DNA rather than RNA, which is the focus of this effort [20]. Metagenomic NGS is particularly attractive for surveillance of febrile illness because the approach can broadly detect viruses, bacteria, and parasites in clinical samples [7, 11]. Although currently limited by a sample-to-answer turnaround time of >24 hours for benchtop sequencers, we report in this study that unbiased pathogen detection using both targeted and WGA NGS which can be realized with actionable results for clinical diagnostics [6, 26, 30, 40] and public health [4, 10, 29] within 10 h.

The characterization of upper respiratory disease (URD) viruses would certainly benefit the use of whole genome sequencing (WGS) and targeted sequencing, providing us with tools for surveillance of highly diverse viral genomes [23, 30]. Such data can assist in vaccine development, detection of anti-viral drug resistance, or the identification of new emerging re-assorted viral genomes. These advances may help to mitigate the morbidity and mortality of influenza pandemics or seasonal viral epidemics. NGS platforms such as the Illumina MiSeq and Life Technologies Ion Torrent have been extensively used to provide WGS data for viruses, potentially within 2-3 days of the receipt of a sample [36]. Furthermore, although the cost of NGS is decreasing, however it is still cost prohibitive unless large numbers of samples are processed in a multiplexing format. Therefore, there is clearly a place for a portable NGS instrument that can be deployed into the field for surveillance using available field ready kits and sequencing reagents that are comparable to cost of benchtop sequencer. Analysis of sequencing data is accomplished in real time by an internet-connected laptop. Another unique feature of the MinION is its ability to generate very long reads of up to 60 kilobases (kb) [28]. The MinION has been successfully used for WGS and targeted sequencing [8, 20, 38].

The purpose of this study is to examine the efficacy of using the MinION to sequence URD organisms present in clinical nasal washes using targeted amplification, WGA and whole transcriptome amplification (WTA) of low titer URD samples.

## Materials and Methods

### Nucleic acid extraction

Influenza A virus H1N1 (VR1736), parainfluenza virus 1-3 (VR94, VR92, and VR93), coronavirus 229E and OC43 isolates (VR-740 and VR1558), *Mycoplasma pneumoniae* (ATCC 29342), *Haemophilus influenzae* virus (51907), and human adenovirus (VR-1603) were obtained from American Type Culture Collection (ATCC) and used as control nucleic acid (NA) for library preparations. RNA/DNA were extracted from 200 μL viral culture supernatants or from clinical nasal washes using the Maxwell total viral nucleic acid kits (Promega), with elution into 50 μL of RT-PCR molecular grade water (Ambion). De-identified positive nasal washes were obtained from clinical laboratory (USAFSAM) included influenza A virus, influenza B virus, parainfluenza viruses 1-3, adenovirus, and coronavirus OC043.

### Targeted RT-PCR and PCR

Standard targeted PCR was done per using primers listed in Table 1 and the SuperScript III OneStep RT-PCR system (Thermo Fisher Scientific) per the manufacturer’s instructions. Briefly, 10 ng of DNA-free RNA or RNA-free DNA was used for each 50-μL reaction. All PCR reactions were accomplished with 1.0 ul of each primer at 10 uM. RT-PCR thermocycling parameters were as follows: 30 minutes at 50°C, 2 minutes at 95°C, and then 35 cycles of 30 seconds at 95°C, 30 seconds at 52°C, and 75 seconds at 68°C, followed with a final extension at 68°C for 5 minutes. PCR thermocycling was accomplished with AccuStartII PCR Supermix (Quanta Biosciences) with following parameters: 3 minutes at 95°C, and then 25 cycles of 30 seconds at 94°C, 30 seconds at 55°C, and 90 seconds at 72°C, followed with a final extension at 68°C for 3 minutes. DNA or cDNA were cleaned up using 1.5X Agencourt Ampure XP beads as described by the manufacturer (Beckman Coulter). DNA and cDNA were quantified using a Qubit 3.0 fluorometer (Life Technologies) and resuspended in Low TE (Invitrogen).

**Table 1.**
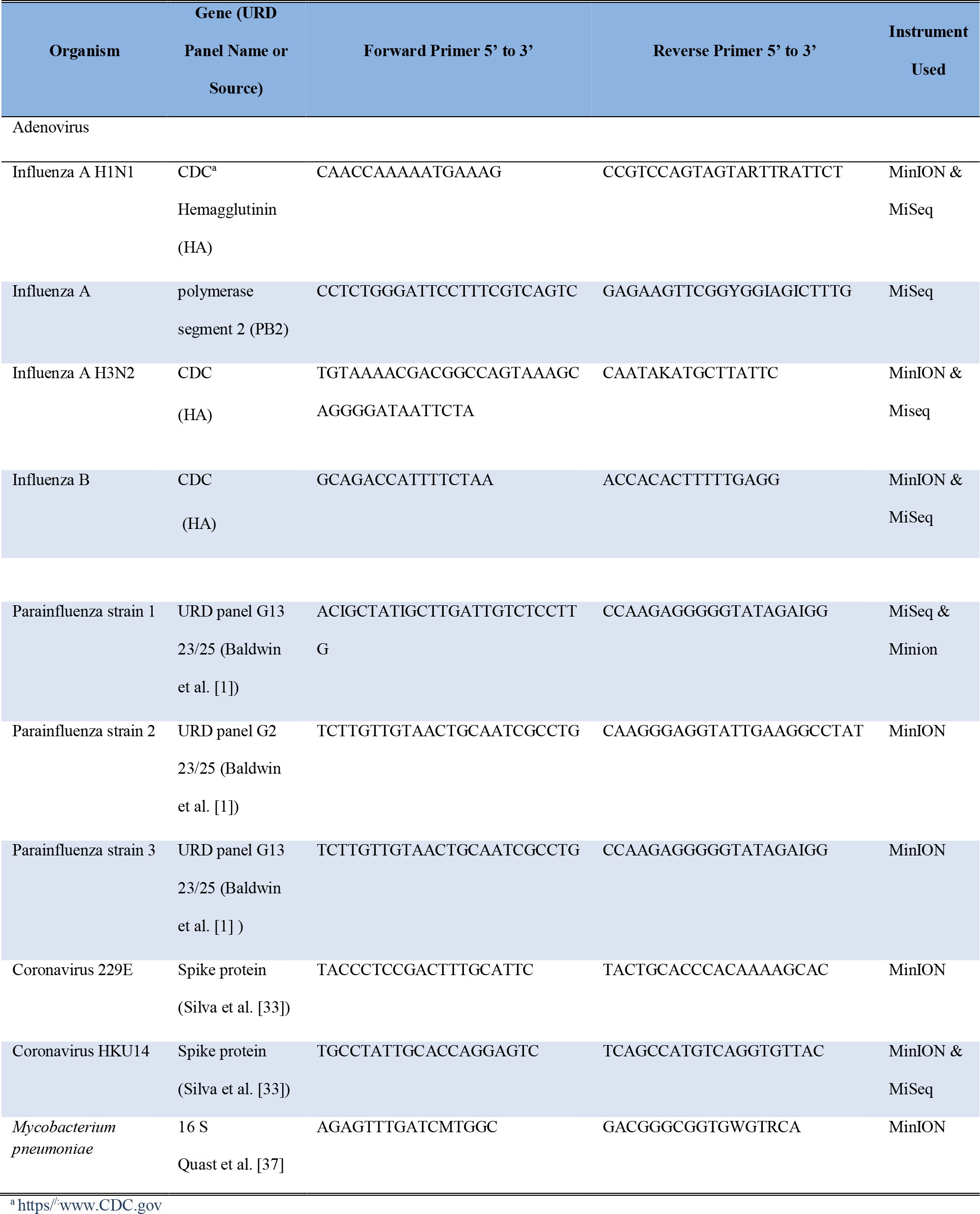
Primers used for targeted RT-PCR.

### WGS preparation

The Oxford Nanopore Technologies (ONT) MinION Genomic DNA Sequencing kit was used to prepare libraries from WGA that previously amplified using REPLI-g WTA/WGA kits (Qiagen) as described by the manufacturer. WTA was used on clinical nasal washes with low viral titers. RT-PCR or PCR was accomplished as described previously. Briefly, 10 ng of DNA-free RNA or RNA-free DNA was used for each WTA/WGA reaction. Whole genome products were sheared with a Covaris g-tube (Woburn, MA) per the instructions of the manufacturer with fragmentation sizing to approximately 5 kb and the sheared DNA was used in library preparations.

### MinION library preparation

This work was completed as part of the ONT MinION early access program. During this time the flow cells, flow cell chemistries and library kits changed numerous times during the two-year study. The ONT MinION Sequencing kit that was used to prepare libraries from WGA/WTA are shown in Table 3. DNA or cDNA libraries were sequenced with various flow cells throughout the study including versions R7, R7.3, and Spot-ON 9.4. The subsequent increase in flow cell number indicates improvements in their flow cells, with Spot-On R9.4 flow cell being the latest improvement. In addition, the ONT MinION library reagents were changed as the kit improvements were made during this study (Kit: SQK-MAP0005 for the R7.03 flow cell and Kits: SQK-LSK- 209, SQK- RAD 004 (Rapid) for R9.4 or Spot-ON R9.4 flow cells). Barcoding of the DNA fragments was completed using barcoding kit (EXP-PBC001). Libraries kit used for each of the sequencing libraries are shown in Table 2 and 3, as well as the amount of starting material and flow cells type. Library preparations were done as described by the manufacturer and were quantified using a Qubit 3.0 fluorometer (Life Technologies).

**Table 2.** Targeted Sequencing of URD samples using the MiSeq and MinION.

| Descriptor | Influenza A<br>H1N1 | Influenza A<br>H1N1 | Influenza B | Influenza B | Parainfluenza<br>1 | Parainfluenza<br>1 | Coronavirus<br>OC043 | Coronavirus<br>OC043 |
| --- | --- | --- | --- | --- | --- | --- | --- | --- |
| Flow cell version | MinION<br>R7.3 | MiSeq<br>600 V3 | MinION<br>R7.3 | MiSeq<br>600 V3 | MinION R7.3 | MiSeq<br>Nano | MinION<br>R7.3 | MiSeq<br>Nano |
| Base calling<br>version/ alignment | Metrichor &<br>BWA/SAM<br>Tools<br>Kraken/<br>Bracken | Illumina<br>Basespace<br>MiSeq/<br>BWA/SAM<br>Tools | Metrichor &<br>BWA/SAM<br>Tools<br>Kraken /<br>Bracken | Illumina<br>Basespace<br>MiSeq/<br>BWA/SAM<br>Tools | Metrichor<br>BWA/SAM<br>Tools<br>Kraken/<br>Bracken | Illumina<br>Basespace<br>MiSeq<br>BWA/SAM<br>Tools | Metrichor<br>BWA/SAM<br>Tools<br>Kraken/<br>Bracken | Illumina<br>Basespace<br>MiSeq/<br>BWA/SAM<br>Tool |
| Library protocol<br>version and<br>amount of starting<br>material | SQK-MAP005<br>1000 ng | Nextera XT<br>1.0 ng | SQK-MAP005<br>1000 ng | Nextera XT<br>1.0 ng | SQK-LSK109<br>1000 ng | Truseq<br>Nano LT<br>100 ng | SQK-LSK109<br>1000 ng | Truseq<br>Nano LT<br>100 ng |
| Target gene | HA | HA | HA | HA | HA | HA | Spike<br>protein | Spike<br>protein |
| PCR amplicon<br>base pair length | 763 | 763 | 886 | 866 | 200 | 150 | 400 | 400 |
| Total number of<br>reads passing QC<br>reads | 114 | 202701 | 7339 | 230049 | 404 | 396725 | 114 | 612294 |
| Total number of<br>aligned reads | 104 | 176744 | 7244 | 212748 | 75 | 365403 | 104 | 597202 |
| 100 base error rate | 1.3E-01 | 3.63E-02 | 1.54E-01 | 7.89E-02 | 1.01E-01 | 2.74E-02 | 1.37E-01 | 3.47E-3 |
| Percent alignment | 87.3 | 97.5 | 98.7 | 93.2 | 18.6 | 92.1 | 91.2 | 97.5 |

**Table 3.**
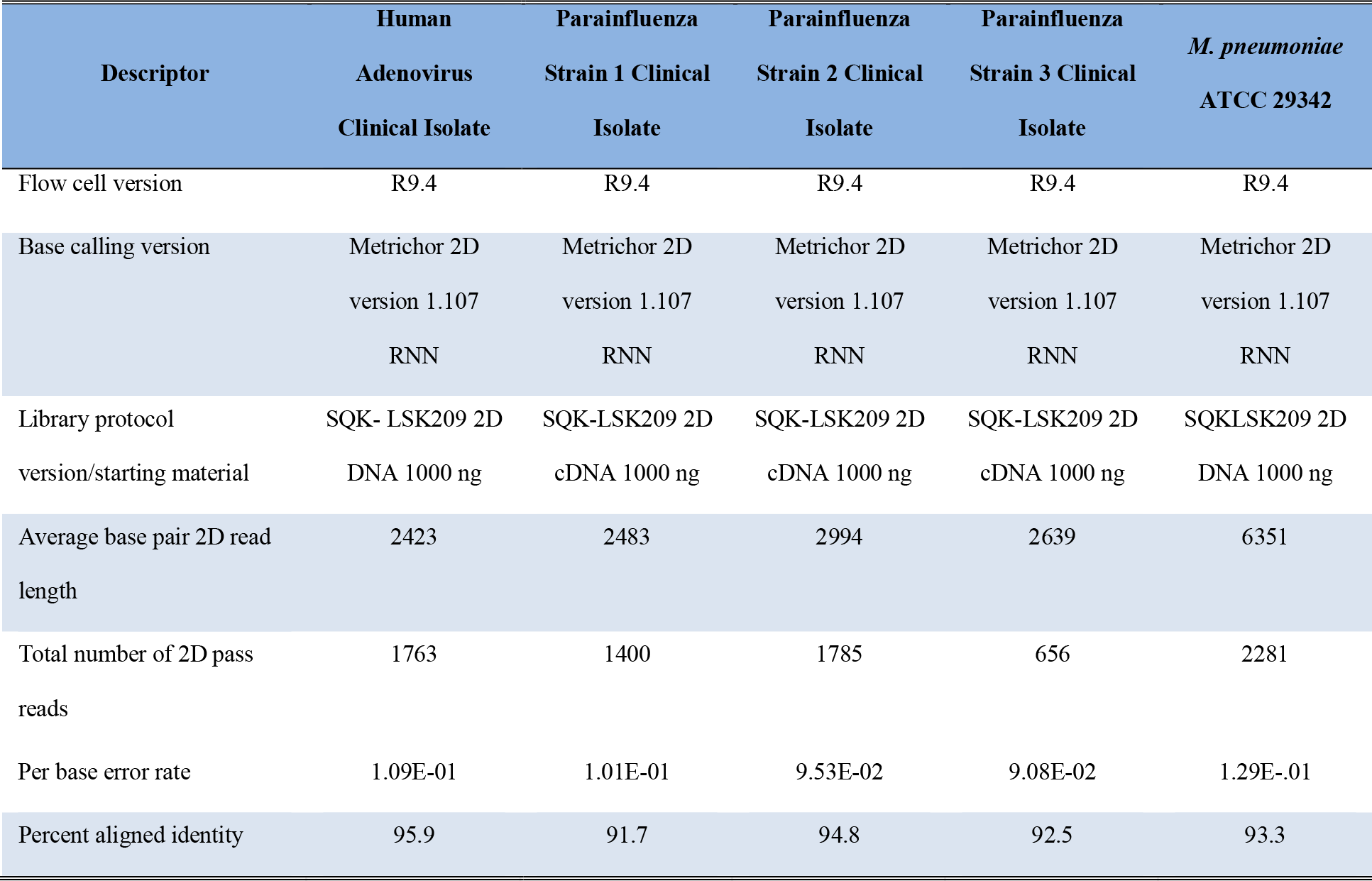
MinION WTA and WGA with Targeted RT-PCR or PCR Amplification.

In each case, the cDNA/DNA was end-repaired in a 100-μL reaction containing 0.5-2 μg of nucleic acid (depending on the run) in 80 μL water, 10 μL of 10X end-repair buffer (from the NEB Next End Repair Module), and 1 μL of enzyme mix. DNA and cDNA were quantified using a Qubit 3.0 fluorometer (Life Technologies) and resuspended in molecular grade water as the starting material for library preparation per the instruction of the manufacturer. End-repair was performed per the manufacturer’s instructions with the exception that the incubation time was increased to 15 minutes. The resulting blunt-ended DNA was cleaned up using 1.0 × volume of AMPure XP beads (Beckman Coulter) according to the manufacturer’s instructions with the exception that 80% ethanol was used instead of 70%, and the DNA was eluted in 30 μL molecular grade water. Ten μL of adapter mix and 2 μL hairpin (HP) adapter were added to the 30 μL dA-tailed DNA, then 8 μL of molecular grade water followed by the addition of 50 μL Blunt/TA ligase master mix (NEB). The contents were briefly mixed and the reaction was left to proceed at room temperature for 15 minutes. One μL of HP tether was added, mixed briefly, and incubated 15 minutes at room temperature. The sample was cleaned up using 1.0 × by volume AMPure XP beads according to the manufacturer’s instructions with the exception that the DNA was eluted into 25 μL of elution library buffer at 37°C for 10 minutes. The beads were removed by placing the tube into a 1.5-mL magnet and the supernatant was transferred to a low binding 1.5-mL Eppendorf tube and placed on ice until the sequencing flow cell was loaded. Each library was quantified using the Qubit fluorometer, and the total library was within the required range of 150-200 ng prior to proceeding to sequencing.

### MinION sequencing

A 48-hour sequencing protocol was initiated using the MinION control software, MinKNOW versions 1.1-1.3.24, for flow cells 7.3 and MinKNOW versions 1.4.6.-2.4.5 for R9 flow cells, respectively. Read event data were base-called by the Metrichor agent (version 0.46.1.9, updated to EPI2ME) using workflows WIMP (What In My Pot) 1.2.2 rev 1.5 with appropriate 2D and 1D workflow scripts [17].

### Illumina MiSeq sequencing

The targeted RT-PCR amplification was accomplished as described earlier using the same RNA as described previously. The sequencing libraries were prepared using either the Illumina TruSeq Nano or Nextera XT library kits (Illumina) with150-bp paired-end sequencing on the Illumina MiSeq.

### Sequence data analysis

Read data were extracted from the native HDF5 format (Fast 5) and converted into FASTA using poretools [25] and sequence reads were assess for quality and trimmed with Nanofilt [9] Quality reads with a q-score lower than 7 were omitted. Illumina reads were trimmed with Trimmomatic 0.36 using the following settings: trim reads: Leading: 3, Trailing: 3, and Sliding Window: 4:15. Taxonomic identification was used to verified for each data set by using bioinformatics packages Burrows-Wheeler Alignment MEM (BWA; aligned to reference and Human GR2) and SAM Tools was used to extract statistics and error rate per 100 bases [21, 22]. Sequence reads were filtered for any human sequence contributions prior to visualization. Tables 2 and 3 listed the analysis tools used for each data set. Kracken and Bracken bioinformatics packages with a k-mer database of viral and bacterial genomes [27, 40]. All genomes, viral and bacterial were downloaded from NCBI Ref Seq (May, 2015) and were used to build the Kraken database (version 0.10.4) with k=31. Bracken was used for reassign alignments to the species level and visualization was accomplished with Krona [31] using the highest of number of hits (lowest e-value) or Metrichor’s WIMP. Viral and bacterial databases were obtained from NCBI viral genomes resource [2]. Read percentage identity is defined as 100 * matches/ (matches + deletions + insertions + mismatches). Fraction of reads aligned are defined as (alignment length + insertions - deletions)/ (alignment length + unaligned length - deletions + insertions) [21].

## Results

### Targeted amplicon sequencing using the ONT MinION

A panel of URD viruses (influenza A virus, influenza B virus, parainfluenza virus 1, parainfluenza virus 2, parainfluenza virus 3, human adenovirus D, coronavirus OC043 and bacterial strains (*M. pneumoniae* and *H. influenzae*) were used in this study. The same nucleic acid from each sample was used to generate libraries for both MinION and Miseq. The MiSeq was used verify the Metrichor WIMP identification. All of the organisms listed above were successfully sequenced on the MinION and identified correctly using the WIMP application, uses a databank of reference sequences for viruses, bacteria, and fungi. Figure 1 shows an a representative example of the data analysis for an influenza A H1N1 isolate listed in Table 1. All viruses and bacteria listed in Tables 2 and 3 were correctly identified using WIMP. In addition, BWA and SAM Tools was used to verify that the alignment for each of the isolates listed in Tables 2.

**Figure 1.**
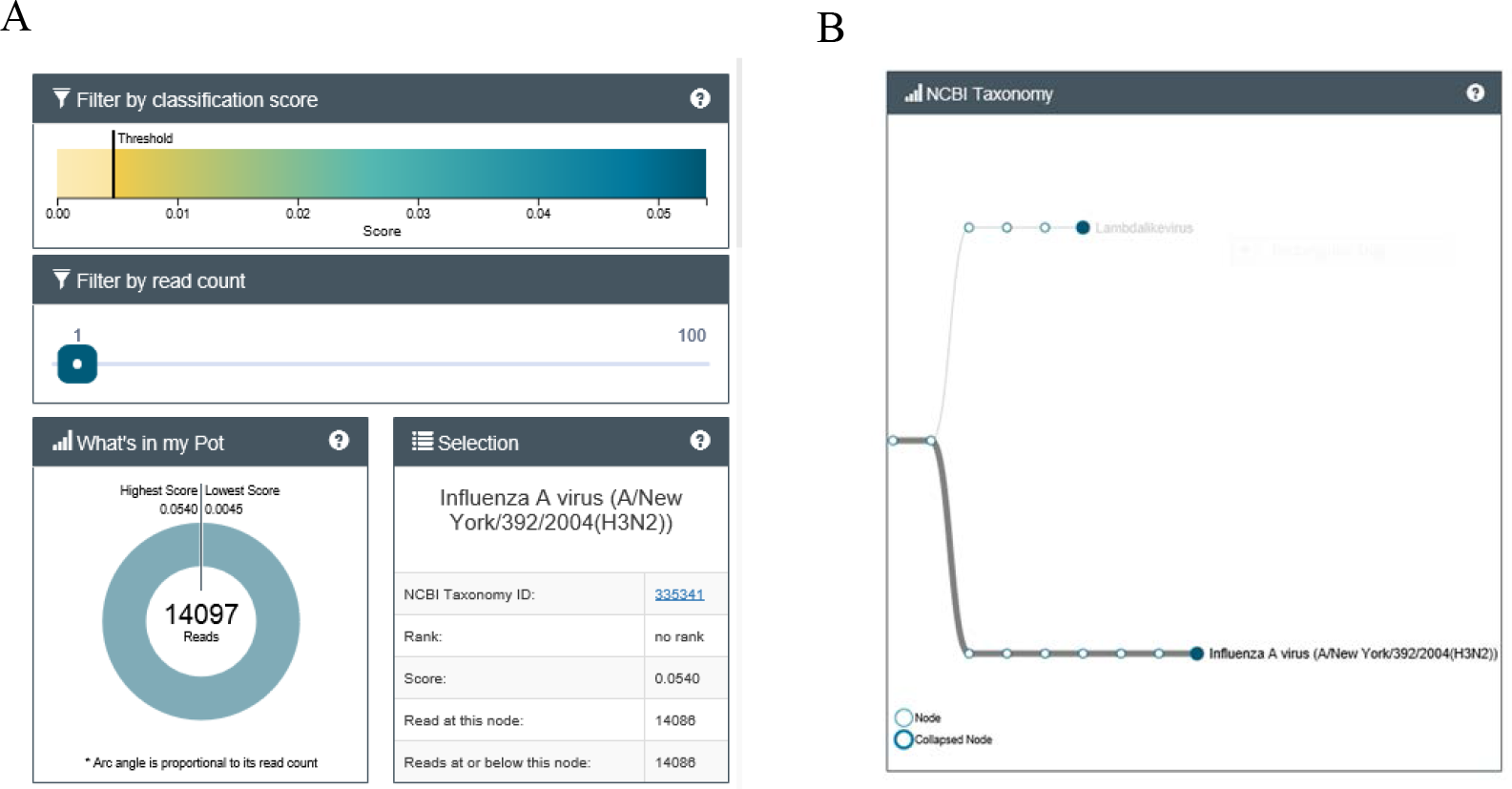
Target amplification of influenza A virus using the H2N2 primers listed in Table 1. Metrichor WIMP application was used for identification of the Influenza A H3N2 virus.

Fast 5 files generated by MinKNOW were converted to FASTQs using Poretools then the Fastq files were aligned with Bracken. For example shown in Table 1, the Influenza A analysis had 114 high quality sequence reads and 104 reads (87.3%) were classified as Influenza A (Table 2).

In addition, MinION sequence identification was further verified by using the same NA extractions for libarary preparation for the Illumina’s Miseq. BWA MEM alignment of the reads were in agreement with Metrichor’s identification for each of the nasal wash samples listed in Table 2. MinION sequence data for viruses Influenza A H1N1, Influenza B, Parainfluenza 1(PAIV1) and Coronavirus OC043 had a percent identity of 82.6, 98.7, 18.4 and 92.4, respectively (Table 2). Interestingly, there was little difference between the sequencing data quality in terms of alignment error. Influenza A H1N1, Influenza B, Parainfluenza 1(PAIV1) and Coronavirus OC043 had alignment error of 1.15E-01, 1.54E-01, 2.74E-02 and 1.37E-01, respectively (Table 2). The Nanopore kits and flow cells used early in this study perform well in regards to identifying the isolates; however, the number of reads that passed were lower than expected probably due improper amplicons to adapter ratios used in the ligation step.

### MinION Sequencing of WGA /WTA amplified NA and targeted PCR Libraries

WTA was used with NA extracted from low titer clinical nasal washes and the isolated RNA (1.0 ng/ul) or DNA (1.0 ng/ul) was amplified with Qiagen’s WTA or WGA kits and the resulting amplification products were used as starting material for RT-PCR or PCR prior to library preparations. The individual libraries were barcoded and mixed in equal molar concentrations prior to being loaded onto a R9.4 flow cell. The percent-alignment to reference genomes ranged from 92-96% (Table 3).

There was minimal sequencing bias seen within the barcoded library pool (data not shown); however, the number of reads were somewhat different among the tested organisms and probably due to an error in quantification of the input library prior to pooling. Future barcoding pooling protocols should include a Tapestation (Agilent) quantification in terms of fragment length, which can be used to calculate of the nM concentration; nevertheless, this experiment demonstrated that the use of Qubit quantifications is sufficient for rapid pooling of multiplex libraries.

Metrichor’s cloud basecalling WIMP application correctly 2D analyzed, demultiplexed sequences and identified the organisms for each virus and bacteria in the library pool (Figure 3F). All of the URD organisms listed in Table 3 were reevaluated using Bracken to verify Metrichor’s identification and visualized with KRONA (Figure 3A-E).

**Figure 2.**
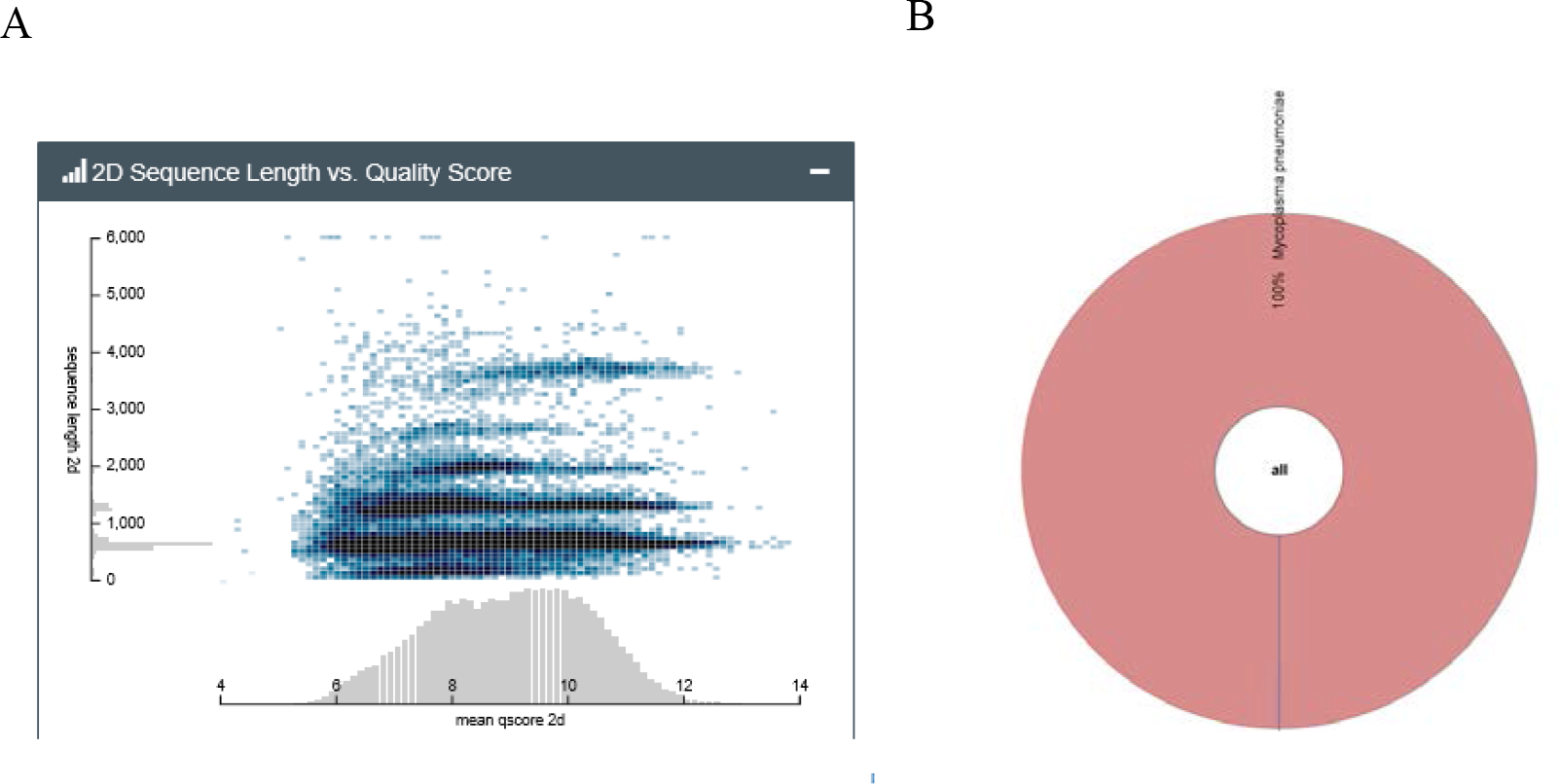
*M. pneumoniae* 2D library identification using Metrichor and Kraken/Bracken. (A) 2D sequence (template + complement strands) as a function of quality score. (B) Bracken was used to align the reads and top hit abundance reads were visualized with KRONA

**Figure 3.**
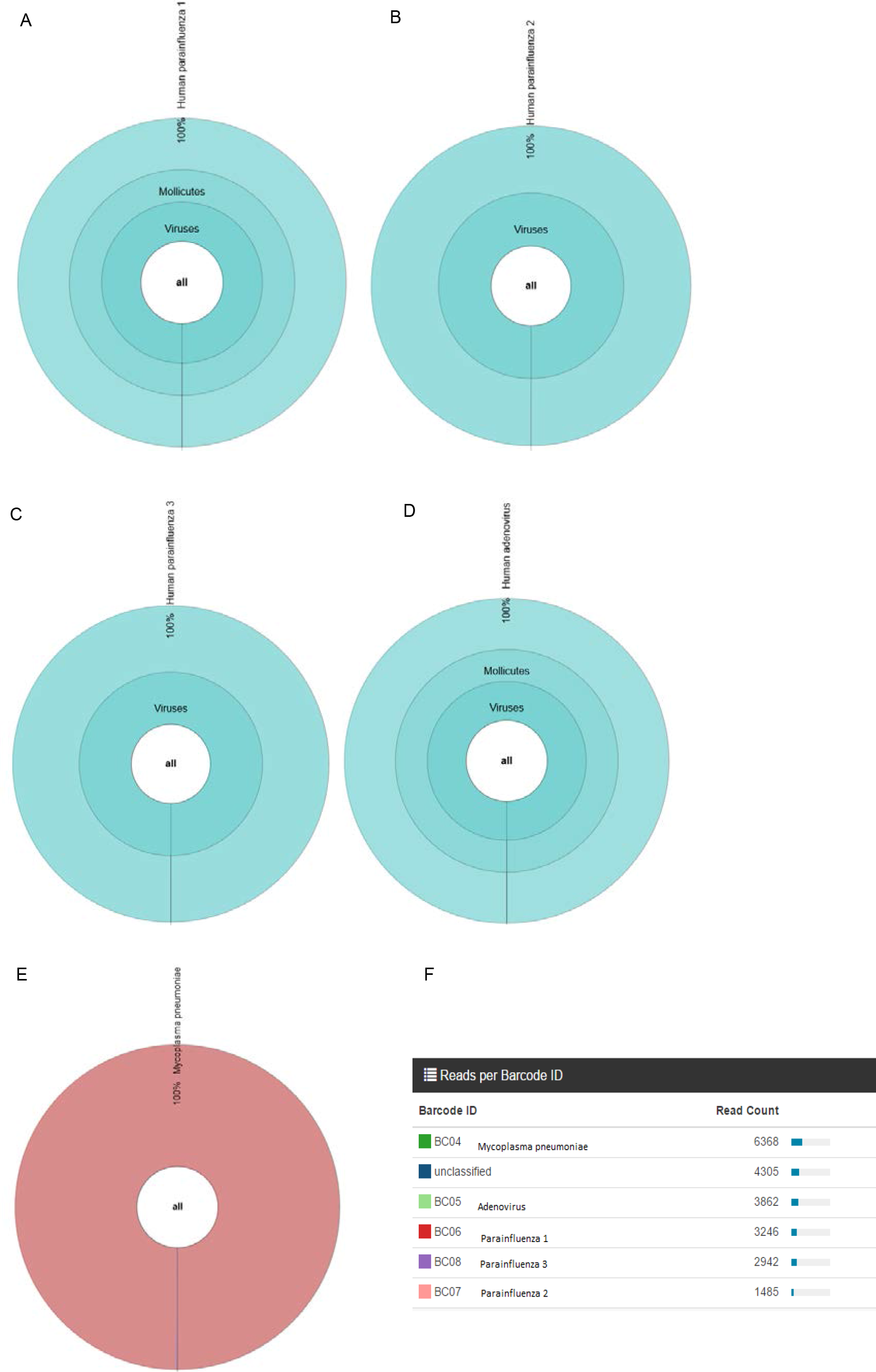
Multiplexing libraries of URD organisms. (A-E) Barcoded URD reads were aligned with Bracken and visualized with KRONA. (F) Number of reads for each barcode after analysis with the Metrichor application 2D base calling.

### 1D Rapid Library Preparation

*H. influenzae* and Coronavirus 229E libraries were prepared using a rapid 1D library preparation. This approach is suitable for WGA products and purified DNA with long fragments. The average Coronavirus 229E length from WTA was approximately 10 kb and the *H. influenzae* was 25 kb prior to library preparation. The *H. influenzae* sequencing run was allowed to run for 17.5 hours The total base yield was 44.85 million and a quality score was centered on 9 (S1). The read length was from 2000 bp and 24050 reads with the majority of the fragments at approximately 5000 bp. *H. influenzae* was correctly identified by the Metrichor’s WIMP (Figure 4A) and the read alignment was verified with Bracken and visualization of the output was accomplished with Krona (Figure 4B). Coronavirus 229E was also successful identified using Metrichor’s WIMP application (S2). The coverage was calculated for both the *H. influenzae* and Coronavirus 229E with the formula C=LN/G, where C = coverage, L = average sequence read length, average Length N = number of reads, and G = the genome size. *H. influenzae* and Coronavirus 229E genome coverage was approximately 186X and 1700X, respectively.

**Figure 4.**
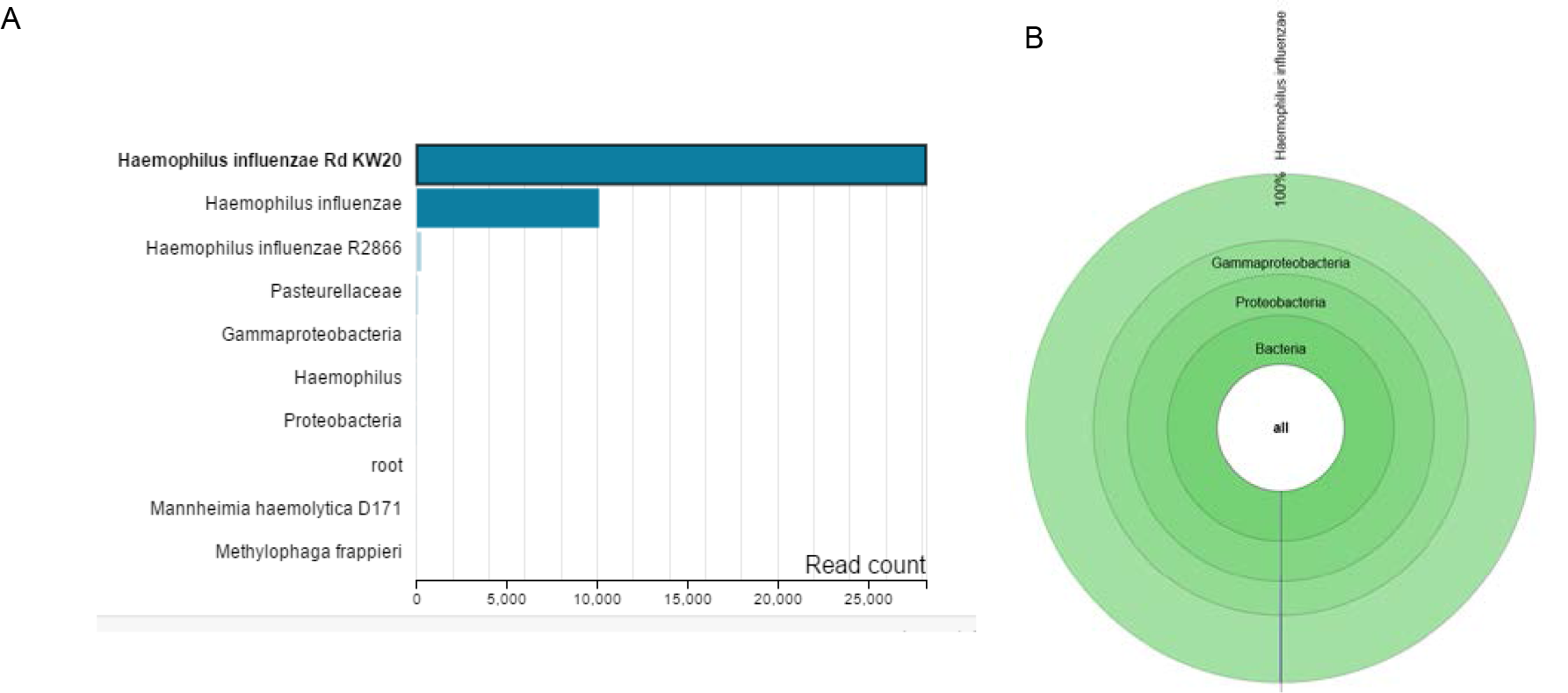
*H. influenzae* was prepared for sequencing using a rapid 1D method. (A*) H. influenzae* read alignment using Metrichor WIMP application. (B) *H. influenzae* reads were aligned with Bracken and visualized with KRONA.

## DISCUSSION

Sequencing of URD viruses and bacteria was accomplished on the MinION nanopore sequencer using both whole genome and amplicon sequencing. Having the ability to sequence amplicons and whole genomes in a mobile, small-footprint platform is attractive when collecting and analyzing samples in the field, and these qualities are also desirable as sequencing methods move toward environmental and surveillance applications. These methods, data, and results represent a practical and novel application for utilizing MinION nanopore sequencing technology. The data in this study were generated with both R7.3 and 9.4 chemistry flow cells, which accurately identified a panel of URD organisms. As the MinION technology improves, sequencing will generally become cheaper, faster, and more accurate as demonstrated in this study with improvements in identity alignment in the >92% range (Tables 2 and 3) as compared to earlier experiments that were in the 30% range [38].

The discrepancies observed between whole genome followed with RT-PCR and amplicon sequencing were minimal in regards to identity alignment (Tables 2 and 3). The base-calling analysis for the MinION nanopore sequencer is an area of active development, as well as flow cell configurations, protocols for kits, and associated kit chemistries. The best quality sequence data are termed 2D reads, while 1D reads are from the template strand. Although the accuracy of MinION sequence data is an area of active development, with recent development of a new nanopore for the flow cells, one should expect increased data quality with each increment in flow cell configuration. Data quality has improved and was demonstrated in this study with each improvement in kit chemistry and flow cell type. This study shows that the platform can be applied for rapid microbial detection with 3-hour run time, which provides sufficient data to generate a reliable identification. As sequencing yield, quality, and turnaround times continue to improve, it is anticipated that nanopore sequencing will challenge mainstays such as PCR for identification of unknown samples. Amplicon and rapid WGS was successfully utilized as an approach for pathogen detection. Low titer viral nasal wash samples were was positively identified after WGA even when samples had high human read counts due to a carry-over human NA contribution. Parainfluenza 1, Parainfluenza 2, Parainfluenza 3 and adenovirus nasal wash samples had 81%, 90%, 95% and 81% of the read count aligning to human references genomes, respectively.

A comparison of the nanopore and short-read sequencing data (Illumina MiSeq) showed that there was agreement in major taxonomic units identified. Therefore, the methodologies described in this study demonstrate that both whole genome and amplicon sequencing will be useful for rapid, accurate, and efficient detection of microbial and viral diversity in clinical samples. In this study, metagenomic sequencing of organisms in clinical samples were successful with increased sequencing depth when WGS was used over that of amplicon sequencing which is beneficial for studying viral genetic rearrangement. In addition, these data also support the utilization of amplicon sequencing over deep sequencing for field applications where surveillance is needed.

The rapid 1D kit, defined as a single strand library composed of template strand, was used to evaluate whole genome amplification (WTA or WGA) of viruses and bacterial species. The library preparation utilizes a transposase to fragment the DNA. Analysis in this study was limited in scope but demonstrates it is functionally a game changer when time is of importance in identifying pathogens in acute infections or during surveillance. This approach is especially beneficial when identification is difficult using conventional bacterial culture techniques and in some cases where organisms are difficult to grow in cultures. Establishing a system for rapid microorganism identification via metagenomic sequencing seems pertinent, especially for field applications.

Although long reads generated from the nanopore sequencer were found to have relatively higher error rates when compared with benchtop sequencing reads. MinION sequencing has been shown to have significant advantages, such as portability, low cost, and real-time detection. These advantages suggest that it is a potential for identification of infectious microbiota community *in situ*. Nevertheless, both whole genome low titer sample such as nasal washes (e.g. WTA, WGA) and amplicon sequencing are more desirable over culturing and will provide a method for identification of organisms in clinical samples. Throughout this study, it has been demonstrated that a low number of long reads are sufficient for accurate identification of small genomes.

## CONCLUSIONS

Recent preliminary reports within the MinION community suggest that the latest version of flow cell chemistry (Spot-On) utilizes more pores per sequence run and generates more data (gigabyte range) when configured with the new nanopore type. Spot-On 9.4 flow cell and flow cell chemistry have improved the amount of errors in sequencing data with reports of >92% sequence alignment and on average <1.2E-01 base error rate per 100 bases. These results confirm the observation by others that the per-base accuracy of MinION data higher that of benchtop DNA sequencing methods (Illumina MiSeq, Table 2 and 3). Targeted sequencing is useful even with higher error rates since gigabytes of data are collected for each run and the fragments are longer (range 800-3 kb) with overlapping sequences which can assist in mitigating error rates.

Having the ability to sequence in a mobile, small-footprint platform is attractive when collecting and analyzing samples in the field, and these qualities are also desirable as sequencing methodology moves toward an even smaller footprint with the ongoing development of a cell phone interface called the SmidgION. These methods, data, and results represent a practical and novel application for the current stage of MinION nanopore sequencing technology and future applications using RNA direct will be a beneficial for rapid identification of RNA viruses. A comparison of the nanopore and short-read sequencing data (Illumina MiSeq) showed that there was agreement in major taxonomic units identified. Therefore, the methodologies described in this study demonstrate that both whole genome and amplicon sequencing will be useful for, accurate, and efficient detection of microbial and viral diversity in clinical samples.

### Availability of supporting data

The raw sequence data sets are available at: https:figshare.com/articles/combined_fastq_zip/7172600

## Acknowledgements

This work was supported by the Defense Health Program and the Chief Scientist Office at Wright Patterson Air Force Base, 711^th^ Human Performance Wing, Ohio. The authors acknowledge the contribution of Mr. Craig Strapple for performing the bioinformatics analyses.

